# Directed disruption of IL2 aggregation sites produces designer biologics with enhanced specificity coupled to improved production capacity

**DOI:** 10.1101/2024.09.02.610760

**Authors:** Amy Dashwood, Ntombizodwa Makuyana, Rob van der Kant, Arman Ghodsinia, Alvaro R. Hernandez, Stephanie Lienart, Oliver Burton, James Dooley, Magda Ali, Lubna Kouser, Francisco Naranjo, Matthew G. Holt, Frederic Rousseau, Joost Schymkowitz, Adrian Liston

## Abstract

The pleotropic nature of interleukin-2 (IL2) has allowed it to be used as both a pro-inflammatory and anti-inflammatory therapeutic agent, through promotion of regulatory T cell (Treg) responses via the IL2RA receptor or promotion of CD8 T cell responses via the IL2RB receptor, respectively. However, the utility of IL2 as a treatment is limited by this same pleiotropy, and protein engineering to bias specificity towards either the regulatory T cell (Treg) or CD8 T cell lineage often requires a trade-off in protein production or total bioactivity. Here we use SolubiS, a computational algorithm-based method, to predict mutations within the IL2 structure to improve protein production yield while altering cellular selectivity, to generate a mutein with elevated therapeutic potential. The design and testing process identified the V126R (murine) / V111R (human) mutation as a Treg-enhancing mutein, creating a cation repulsion to inhibit primary binding to IL2RB, with a post-IL2RA confirmational shift enabling secondary IL2RB binding, and hence allowing the trimeric receptor complex to form. In human IL2, additional N110R T151R aggregation-protecting mutations could improve protein yield of the V111R mutation. The approach also generated novel CD8 T cell-promoting mutations. Y79K created a cation-cation repulsion with IL2RA, while Q50W enhanced CD8 T cell activity through potential π-stacking enhancing binding to IL2RB, with the combination highly stimulatory for CD8 T cells. For human IL2, Y65K (homolog to murine Y79K) coupled with E82K prevented IL2RA binding, however it required the aggregation-protecting mutations of N110R T151R to rescue production. These muteins, designed with both cellular specificity and protein production features, have potential as both biological tools and therapeutics.

## Introduction

Interleukin 2 (IL2) is a highly potent cytokine, capable of both maintaining homeostasis and driving inflammatory reactions [1].These dual functions are based on the differential expression of the different receptor complexes. The high affinity receptor, a trimer of IL2RA, IL2RB and ILR2G (CD25, CD122, CD132, respectively), is dominantly expressed by regulatory T cells (Tregs), allowing IL2 to control Treg homeostasis through effects on Treg “fitness” and proliferation/apoptosis control [2] [3, 4] [5]. The intermediate affinity receptor, a dimer of IL2RB and ILR2G, is expressed by cells with more inflammatory potential, largely CD8 T cells and NK cells. Due to the ∼100-fold lower affinity of the dimer for IL2, at normal levels of IL2 production the dominant role is to establish homeostasis [6, 7]. Elevated IL2 production is, however, among the earliest events following CD4 T cell activation [8], supporting the proliferation of CD8 T cells and NK cells during an immune reaction. This response is aided by transient low-level upregulation of IL2RA on responding cells, providing an inflammatory interlude before homeostasis is re-established through the Treg response to elevated IL2 [7]. This dominant IL2 network is further complicated by context-dependent IL2 circuits [9] and alterations of receptor expression during pathology [10, 11].

With the immunological capacity to both elevate tolerogenic (Treg) and immunogenic (CD8 T cell/NK) responses, IL2 is a highly attractive therapeutic target. The clinical utility is, however, limited by the concentration-dependent effects, especially as the half-life of IL2 is around 5 minutes following injection [12, 13]. Multiple approaches to engineer more stable concentrations have been developed, including modification of IL2 through post-translational modification, such as PEGylation [14], via fusion to protein carriers, such as the immunoglobulin Fc region [15] [16] or a soluble version of the IL2RA protein [17] [18], or through sustained production using gene therapy vectors [19]. An alternative approach has been to decouple this concentration-dependent effect by restricting function to either the trimeric or dimeric receptor, through either screening/*in vitro* evolution approaches [20] or rational design [21] [22]. IL2 muteins can be created with alterations to the IL2RA, IL2RB or IL2RG binding interfaces, including enhanced binding to IL2RA [23], reduced binding to IL2RA [24], increased binding to IL2RB [20, 21, 25]; reduced IL2RB binding [26] or reduced activation of the receptor complex via IL2RG [27] [25], [28], [29], [22]. The net effects are designer IL2 variants with increased selectivity for Tregs, increased selectivity for CD8 T cells and NK cells, or antagonistic activity, depending on which binding structure is disrupted. IL2 variants combining both elevated longevity and altered specificity have also been generated, such as combining an IL2 mutein with an Fc fusion [30], or targeting PEGylation to a particular IL2 interface [31]. Finally, more exotic approaches have been used, including the *de novo* design of protein capable of binding only one surface of the IL2R [32], or the creation of IL2/IL2R orthogonal pairs that do not cross-react with the native system [33].

While IL2 muteins provide enhanced specificity for either the Treg or CD8 T cell lineage, depending on the mutation design, a feature commonly observed in muteins is reduced stability or bioactivity [26], [34], [35], [36]. As either of these attributes limit the utility of the mutein in biotechnology, we sought to design novel IL2 mutations using the SolubiS algorithm, which enables rational design of mutations to enhance stability. Using this system, we designed and validated families of IL2 mutations with enhanced specificity for either Tregs or CD8 T cells, while maintaining total bioactivity and maintaining or even improving production capacity. This combination of features provides potential utility for these muteins in clinical therapeutics.

## Methods

### IL2 mutation design

Mutations were designed based on four structures of human IL2: 1m47 (IL2 alone, resolution of 1.99 Å [37]), 1z92 (IL2 with IL2RA, resolution of 2.8 Å [38]), 2b5i (IL2 with IL2RA/IL2RB/IL2RC, resolution of 2.3 Å [39]) and 2erj (IL2 with IL2RA/IL2RB/IL2RC, resolution of 3 Å [40]), and one structure of mouse IL2: 4yqx (IL2 bound to anti-IL2 single chain Fv, resolution of 2.8 Å [16]). Mutation design was based on the SolubiS method [41] [42] [43] to predict aggregation-prone regions from the primary peptide sequence. The thermodynamic contribution of these regions to the stability of the protein was predicted using FoldX [44], an empirical force field developed for the rapid evaluation of the effect of mutations, and TANGO, a statistical thermodynamics algorithm [42].TANGO calculates the intrinsic aggregation propensity of aggregation-prone regions using a Boltzmann distribution with competing secondary structural tendencies, such as α-helical or β-hairpin structure. The benefit of this implementation is that TANGO predicts aggregation-prone regions [41] with well-defined sequence boundaries, with high specificity and thus predicts few false positives [45]. Each amino acid of the wildtype IL2 protein was mutated to every other amino acid in each structure. The effect of both the thermodynamic stability (ddG) as well as the stability of the interaction (ddGComplex) with each of the receptors was calculated using FoldX (version 3.0 beta 6), and the effect of each mutation on the aggregation propensity was calculated using TANGO, based on sequence alone. This analysis resulted in a large table of FoldX energies and aggregation propensity (TANGO) scores. Mutations were selected using four strategies: SolubiS (reducing aggregation while retaining stability), kill IL2RA binding (reducing interaction energy with IL2RA, while retaining stability and preferably reducing aggregation), kill IL2RB binding (reducing interaction energy with IL2RB, while retaining stability and preferably reducing aggregation), and combinations (identifying synergies that combine aggregation reduction and IL2RA/IL2RB binding). The thresholds used were reduction of TANGO score by more than 100, for reduction in aggregation, while not destabilising hIL2 thermodynamically (ddG < 0.5 kcal/mol) and not affect any receptor interactions negatively (ddG_complex < 0.5 kcal/mol). These single point mutations were calculated in combination to allow for the previously described synergistic effect of SolubiS mutations on the aggregation resistance of proteins [46]. These mutation combinations were selected in the same way as single SolubiS mutations, to ensure compatibility.

Structural models of IL2 in complex with the trimeric receptor were based off the 2erj structure [40] of the human IL2 / IL2R complex. Due to the high homology between mouse and human IL2 and IL2R complex, and the lack of a mouse structure, selected human mutations were directly imposed on the mouse sequence. Each introduced amino acid was the same for both human and mouse mutant. This resulted in some cases where a different wildtype amino acid was mutated to the same mutant amino acid. For example, in the case of the H36W mutation, where the human homologue has a histidine residue, the mouse homologue has a glutamine residue, making the mouse equivalent mutant Q50W. FoldX calculations were performed on the monomeric crystal structure 4yqx (mIL2 alone, resolution 2.8 Å [16]).

### Cloning and molecular biology

IL2 muteins were ordered as Gene Fragments from GeneWiz, Azenta. Gene Fragments were cloned into pJet Blunt 1.2 (ThermoFisher, K1231) and sequenced via Sanger Sequencing (Genewiz, Azenta) to confirm correct sequence insertion. Correct insertions were amplified via PCR adding restriction enzyme sites to each end. PCR products were purified using GeneJET PCR purification kit (ThermoFisher, K0701) before being digested with Pfl23II (ThermoFisher, FD0854) and NdeI (ThermoFisher, FD0583) and gel purified (GeneJET Gel Extraction Kit, ThermoFisher, K0691) to obtain the IL2 sequence. For low-expressing murine muteins V126R, N123H T166R and Y79K E96K T166R, codon-optimised sequences were used. Fragments were then cloned into an expression vector, downstream of an EF-1α promoter and in frame with a TEV protease target and a 6xHis-Tag. Finally, sequences were confirmed using Sanger sequencing (Genewiz, Azenta).

### IL2 production and measurement

Expression plasmids were transfected into HEK293 cells, seeded at 100,000 cells/well, using FuGENE HD Transfection reagent (Promega, E2311) in 24 well plates containing 500µL DMEM supplemented with 10% FBS. 24 hours following transfection, wells were topped with an additional 250µL of DMEM, 10% FBS. After a total of 48 hours, the culture supernatant containing IL2 protein was harvested and frozen at -80°C. All supernatants were aliquoted prior to freezing to prevent multiple freeze thaw cycles. The cells were washed with PBS and detached, and cell pellets were frozen at -80°C ready for DNA extraction. Tranduction efficiency was calculated through PCR quantification of the vector. DNA was extracted from frozen cell pellets using the DNeasy Blood and Tissue Kit (Qiagen, 69504) following manufacturer’s instructions. DNA samples were analysed using QuantStudio 1 via qPCR using TaqMan Gene Expression Assays (FAM) for the detection of plasmid DNA (using the neomycin resistance cassette) relative to genomic DNA detection of GAPDH (Hs02786624_g1) and PPIA (Hs04194521_s1), enabling the transfection efficiency to be calculation for production normalisation.IL2 production was analysed via a modified sandwich ELISA. Plates were coated with anti-IL2 antibodies ((mouse: clone JES6-1A12, BioLegend, 503701) (human: polyclonal, Bio-Techne, AF-202-NA)). The detection antibody was HRP-conjugated anti-Histag (BioLegend, 652503). Standard curves were created using his-tagged IL2 (GeneTex: Mouse; GTX00285-pro, Human; GTX00092-pro), from which concentrations of IL2 in cell culture supernatant were interpolated. To compare production efficiency of each IL2 mutein, final concentrations were normalised by transfection efficiency values calculated from aforementioned qPCR.

### Phospho- STAT Flow cytometry

For murine phospho-STAT assays, spleens dissected from C57Bl/6 mice were mechanically disrupted between two glass slides, filtered through 100µm mesh and red blood cells lysed to form single cell suspensions. Cell suspensions were counted using a CellDrop cell counter. Non-specific binding was blocked using 2.4G2 supernatant. Surface staining was performed with antibodies against CD4, CD127, CD62L, CD25, CD44 as well as an e780 fixable Live-Dead stain (eBioscience). Splenocytes were then stimulated with wildtype IL2 and mutated IL2 diluted to a range of concentrations (400-0.02ng/ml) in complete DMEM. Immediately following stimulation, cells were fixed with paraformaldehyde followed by methanol. An overnight stain [47] was then preformed with antibodies against Foxp3, CD3, CD8, pSTAT5, CD122, CD132 and pSTAT3. Following staining, data was acquired on a Cytek Aurora, with data collection performed in SpectroFlo (Cytek) and analysed using FlowJo and GraphPad Prism.

For human phospho-STAT assays, PBMCs were isolated from leukocytes cones from healthy donors (NHSBT NCI) using Ficoll, following the manufacturer’s instructions. Cell suspensions were counted using a CellDrop cell counter. Non-specific binding was blocked using FcX Trustain (1:200) (BioLegend). Surface staining was performed with antibodies against CD4, CD3, CD8, CCR7, CD45RA, CD27, CD56 as well as an e780 fixable live dead stain (eBioscience). Cells were then stimulated with wildtype and mutant IL2, diluted to a range of concentrations (400-0.02ng/mL) in complete DMEM. Immediately following stimulation, cells were fixed with paraformaldehyde followed by methanol. An overnight stain [47] was the performed with antibodies against FOXP3, CD25, and pSTAT5. Following staining, data was acquired on a Cytek Aurora, with data collection performed in SpectroFlo (Cytek) and analysed using FlowJo and GraphPad Prism.

### In vivo gene delivery and assessment

For gene therapy studies using adeno-associated virus-based vectors (AAVs) male and female C57BL/6 wildtype mice were obtained from Babraham Institute Biological Support Unit. All mice were maintained in a specific pathogen free environment, with ambient temperature, humidity, and lighting maintained for their well-being. Before experimentation, mice were transferred to individually-ventilated cages, and grouped matching age and sex. Mice were fed with either wet or dry standard food pellets based on the animal food consumption preferences.

Wildtype (C57BL/6) mice were intranasally inoculated with 50μL of 1x10^11^ GC/ml of AAVs diluted in PBS. All mouse experimentation and procedures compiled with the UK Home Office guidelines observing the Animals Scientific Procedures Act 1986. Experiments were also approved by the Babraham Institute Animal Welfare and Ethical Review Body (Protocol study number PP3981824-Lymphocyte development and immune function in the tissues).

rAAV constructs were created using AAV6.2 capsid and Scgb1a1 promoter. The expressed gene was GFP-Myc-Cre (Addgene #49056), murine IL2 (NM_008366.3) or murine IL2 V126R (GTA to AGA in codon 126). Vectors were produced by Vector Builder (sterile, <90% purity).

At the indicated time points post-treatment, mice were injected intravenously with anti-CD45 antibody before euthanasia. The lung was harvested into PBS with 2.5% FCS and 2mM EDTA. Tissues were homogenized in Protein Quant Sample Lysis kit Thermo Scientific™ buffer and a protease inhibitor (Complete Mini EDTA Sigma). The resultant supernatant was checked for IL2 levels using ProQuantum High-Sensitivity Immunoassays Kit from Thermo Fisher following the manufacturer’s protocol. IL2 standard curves were done on the ProQuantum platform (Thermo Scientific).

For flow cytometry analysis, tissues were processed as previously described [48]. Samples were prepared by initially blocking Fc interactions with supernatant from 2.4 G2 hybridoma cell lines, followed by viability staining, staining with the surface marker cocktail for 2 hours, and staining overnight for intracellular markers [47] (**Supplementary Table 1**). 10,000 Precision Count beads (BioLegend) were added to samples before running through a Cytek™ Aurora (5 Lasers N9-20001_Spectral). Data analysis, including flow cytometry gating, was performed with FlowJo v.10.0.7 and FlowJo version 10.8.1 (BD).

## Results

### Directed disruption of IL2 aggregation sites designed by SolubiS

A key limitation in the production capacity of proteins is the aggregation of nascent polypeptides, mediated through aggregation-prone hydrophobic stretches [49]. As protein-protein binding sites are frequently hydrophobic, this feature allows the potential for the design of mutations that both disrupt a protein-protein interaction and also improve protein production through lowering the aggregation rates [50]. In order to improve the properties of IL2, we designed a series of mutations in both human and mouse IL2. Mutation design was based on the structure of the human and murine IL2 structures, and used the SolubiS method [46] to provide maximal improvements with minimal point mutations. Candidate mutations were selected for the ability to reduce the interaction energy of IL2 for IL2RA or IL2RB, while reducing the aggregation propensity of the polypeptide. (**Table 1**). A full list of the predicted biochemical properties of each mutein is listed in **Supplementary Spreadsheet 1**.

**Table 1.** IL2 muteins design strategy.

| Organism | Mutation | # of mutations | Human homolog | Strategy |
| --- | --- | --- | --- | --- |
| Human | N110R | 1 | - | SolubiS monomer |
| Human | N110E | 1 | - | SolubiS monomer |
| Human | T143N | 1 | - | SolubiS monomer |
| Human | T151R | 1 | - | SolubiS monomer |
| Human | H36W | 1 | - | Kill IL2RB binding |
| Human | D40R | 1 | - | Kill IL2RB binding |
| Human | N108H | 1 | - | Kill IL2RB binding |
| Human | V111R | 1 | - | Kill IL2RB binding |
| Human | N110R T143N | 2 | - | SolubiS combo |
| Human | N110R T151R | 2 | - | SolubiS combo |
| Human | V111R T143N | 2 | - | SolubiS with kill IL2RB binding |
| Human | N108H T151R | 2 | - | SolubiS with kill IL2RB binding |
| Human | E82K | 1 | - | Kill IL2RA binding |
| Human | Y65K | 1 | - | Kill IL2RA binding |
| Human | Y65K E82K | 2 | - | Kill IL2RA binding |
| Human | Y65K E82K N110E | 3 | - | SolubiS with kill IL2RA binding |
| Human | Y65K E82K N110R T151R | 4 | - | SolubiS with kill IL2RA binding |
| Mouse | A158N | 1 | T143N | SolubiS monomer |
| Mouse | T166R | 1 | T151R | SolubiS monomer |
| Mouse | Q50W | 1 | H36W | Kill IL2RB binding |
| Mouse | D54R | 1 | D40R | Kill IL2RB binding |
| Mouse | N123H | 1 | N108H | Kill IL2RB binding |
| Mouse | V126R | 1 | V111R | Kill IL2RB binding |
| Mouse | V126R A158N | 2 | V111R T143N | SolubiS with kill IL2RB binding |
| Mouse | N123H T166R | 2 | N108H T151R | SolubiS with kill IL2RB binding |
| Mouse | E96K | 1 | E82K | Kill IL2RA binding |
| Mouse | Y79K | 1 | Y65K | Kill IL2RA binding |
| Mouse | Y79K E96K | 2 | Y65K E82K | Kill IL2RA binding |
| Mouse | Y79K E96K A158N | 3 | Y65K E82K N110E | SolubiS with kill IL2RA binding |
| Mouse | Y79K E96K T166R | 3 | Y65K E82K N110R T151R | SolubiS with kill IL2RA binding |

### Production screening of IL2 muteins identifies point mutations with improved production capacity

To functionally test the designed IL2 mutations, we first screened for production capacity of each mutein, through cloning the IL2 muteins into an expression plasmid transfected into HEK293 cells Through normalisation of IL2 protein production (measured through ELISA for a His tag fused to the C terminus of each protein) to HEK293 transfection efficiency (measured through PCR quantification of expression plasmid), we could compare the production output of each mutein relative to wildtype IL2. For the mutations in murine IL2, muteins T116R, V126R A158N, and Y79K resulted in a trend towards elevated IL2 production, up to ∼3-fold higher than wildtype IL2 levels, while other mutants gave equivalent or worse production (**Figure 1A**). For the mutation in human IL2, muteins N110R T143N, V111R T143N trended towards increased production but no mutants gave significantly elevated production and several had production defects (**Figure 1B**). The combination of N110R T151R with Y65K E82K resulted in a rescue of production (p=0.055) equivalent to wildtype levels, whilst the Y65K E82K mutation on its own resulted in a markedly reduced production (**Figure 1B**).

**Figure 1.**
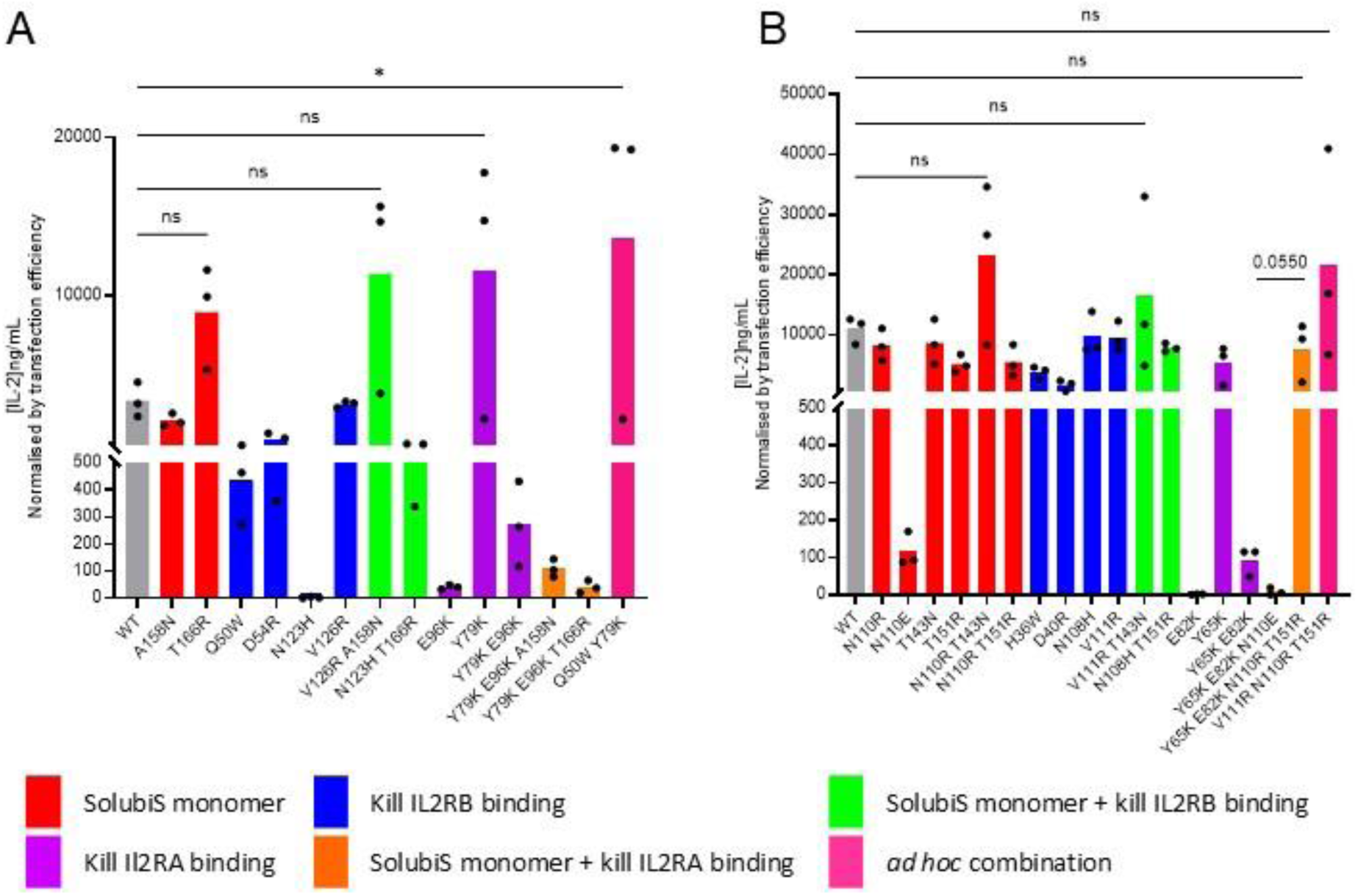
Screening of IL2 muteins identifies muteins with increased production capacity. **A)** Murine IL2 mutein production in mammalian HEK293 cells detected via HisTag ELISA and normalised to transfection efficiency measured by qPCR (n=3). **B)** Human IL2 mutein production in mammalian expression system HEK293 detected via HisTag ELISA and normalised to transfection efficiency measured by qPCR (n=3). Comparisons made using One-Way ANOVA (*P < 0.05, **P < 0.01, ***P < 0.001, **** P<0.0001).

### SolubiS murine IL2 muteins have altered cellular specificity

To determine the impact of these IL2 mutations on receptor specificity, we used a functional assay based on detection of signalling within murine Tregs or CD8 T cells. Using phospho-STAT5 (pSTAT5) as the readout for receptor triggering in both Tregs and CD8 T cells, following incubation in a mixed murine splenocyte population, activation was assessed by flow cytometry. Of the murine mutations designed to prevent binding to IL2RA (E96K and Y79K), both demonstrated reduced pSTAT5 activation in Tregs alone and in combination (**Figure 2A**). In the case of Y79K, this indicated reduced bioactivity on Tregs. E96K, by contrast, also had production issues and was, therefore, tested at suboptimal doses (0.3ng/ml), with the poorer pSTAT5 activation potentially reflecting this fact. Of the murine mutations designed to prevent binding to IL2RB (Q50W, D54R, N123H, V126R), Q50W and V126R had preserved activity on Tregs, while D54R and N123H had reduced activity on Tregs (**Figure 2A**), indicating the latter two mutations had off-target detrimental effects (although in the case of N123H, this may alternatively be attributed to suboptimal production (5.9ng/ml)). For the effects on CD8 T cells, V126R, V126R A158N and N123H T166R combined normal Treg stimulation with the predicted loss-of-signalling on CD8 T cells, while Q50W had an unexpected major gain-of-signalling effect on CD8 T cells (**Figure 2B**). The Y79K mutation retained CD8 T cell stimulation, while having impaired Treg stimulation (**Figure 2A,B**), consistent with impeded IL2RA binding. Residues designed to boost production, A158N and T166R, were neutral to cell selectivity (**Figure 2A,B**).

**Figure 2.**
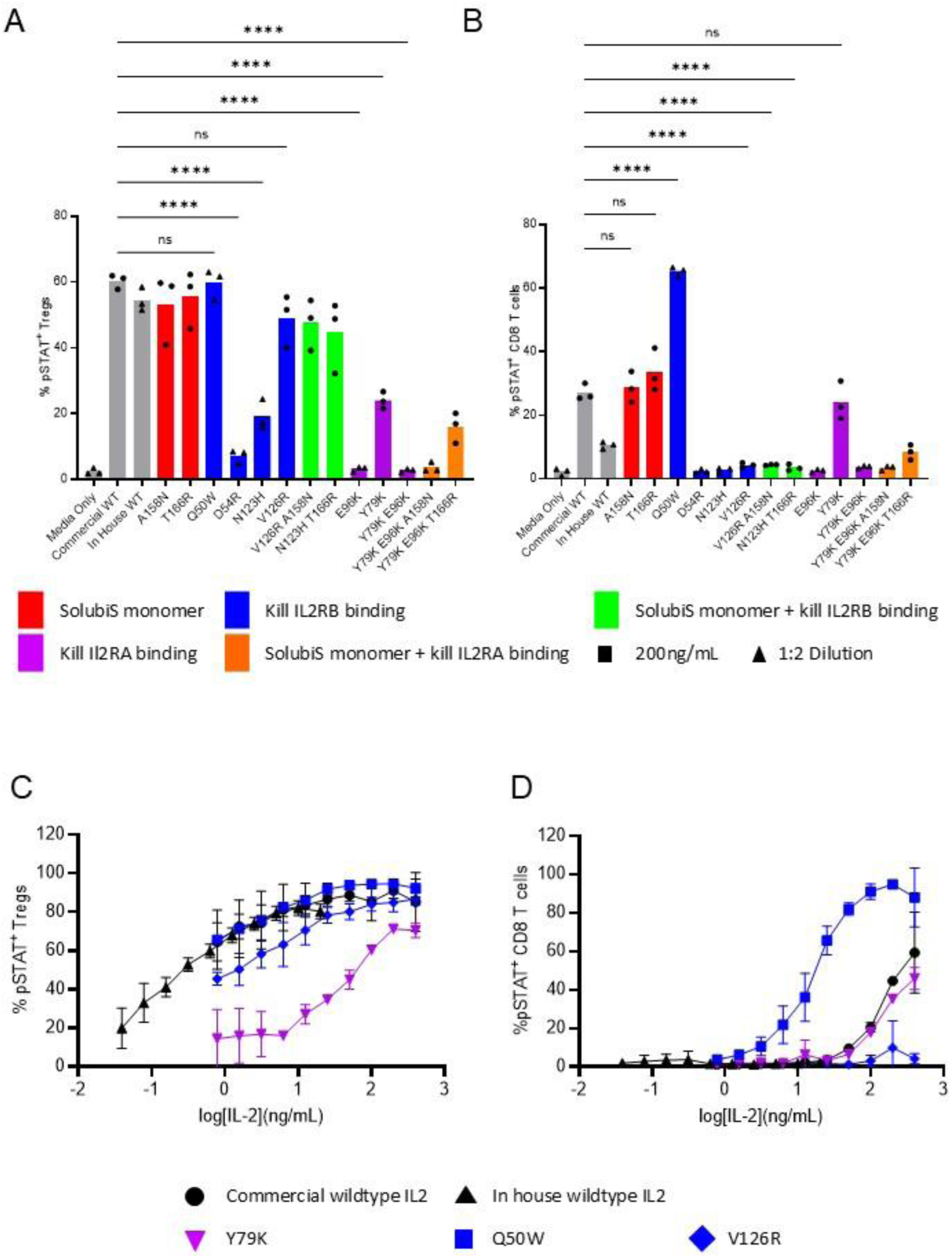
Altered cellular responses to murine SolubiS IL2 muteins. **A)** Mouse splenocytes were incubated for 20 minutes with either media containing 200ng/mL IL2 or 200ng/mL murine mutein, prior to staining for pSTAT2. For muteins with poor production, where ELISA detected less than 200ng/ml, a 1:2 dilution of the production supernatant was used instead. The percentage of pSTAT5+ cells within the Treg or **B)** CD8 T cell populations (n=3). Comparisons made using One-Way ANOVA (*P < 0.05, **P < 0.01, ***P < 0.001, **** P<0.0001). **C)** Mouse splenocytes were incubated for 20 minutes with either media containing IL2 or murine mutein, using a serial dilution of IL2 concentration, prior to staining for pSTAT5. The percentage of pSTAT5^+^ cells within the Treg or **D)** CD8 T cell populations (n=3).

From the initial screen, three murine mutations warranted detailed functional analysis: Y79K, V126R and Q50W. To determine the relative bioactivity of these muteins on Tregs and CD8 T cells, we used the same pSTAT5 response assay, across a titrated dose range, providing enhanced sensitivity for detection of functional changes. Compared to both commercial IL2 and the wildtype IL2 control, Y79K demonstrated a >300 fold reduction in activity on Tregs (**Figure 2C**), while maintaining normal activity on CD8 T cells (**Figure 2D**), consistent with the designed defect in IL2RA binding. The reverse was observed for V126R (designed for defective IL2RB binding), with normal activity on Tregs (**Figure 2C**), but no detectable effect on CD8 T cells, even at high doses (**Figure 2D**). As identified in the functional screening assay, Q50W demonstrated an unexpected effect: normal activity on Tregs (**Figure 2C**), but strongly elevated effects (∼100-fold increased activity) on CD8 T cells (**Figure 2D**). Together, these results indicate Y79K results in CD8 T cell-specific signalling, V126R results in Treg-specific signalling, and Q50W acts as an unexpected CD8 T cell superkine.

In order to identify the potential mechanism by which Q50W displayed an unexpected increase in activity on CD8 T cells, we modelled the effects of the three mutations on murine IL2 / IL2R binding, using a structure built on the human homologs (**Figure 3A**). First, for Y79K, when modelled as bound to the trimeric IL2R, the native Y79 residue of IL2 is in close proximity to positive charges on the R35 and R36 residues of IL2RA (**Figure 3B**), consistent with a cation–π interaction. The replacement of this residue by Y79K creates a cation-cation repulsion, consistent with a loss of binding to IL2RA. For the V126R mutein, as the IL2RB interface of IL2 changes conformation after it binds IL2RA, we modelled the complex in the presence and absence of IL2RA. In the unbound state of IL2, the V126R residue would occupy the same space as R41/R42 on IL2RB, creating a direct cation repulsion effect (**Figure 3C**). However, following binding to IL2RA, IL2 changes conformation in the helix C on which V126R lies, a conformational change that was previously postulated to prime recruitment of IL2RB into the trimeric receptor complex [39]. This conformational change pulls back the V126R residue, reducing the direct cation-cation clash (**Figure 3C**). This is consistent with the observed poor activation through the dimeric IL2RB/IL2RG complex, but unimpeded activation through the trimeric IL2RA/IL2RB/IL2G complex. Finally, the Q50W mutation was modelled as the analogue to the human H36W, based on the human IL2/IL2R structure. While the H36W mutation was predicted to disrupt the interaction, based on the crystal structure of the complex, Q50W was initially only checked for its effect on the stability of the monomer. Subsequent modelling of the mouse complex revealed the Q50W side chain falls within a hydrophobic groove of IL2RB with potential π-stacking interactions with F101 and F135 residues (**Figure 3D**). The larger hydrophobic side-chain may, therefore, enhance binding, providing preferred signalling through IL2RB.

**Figure 3.**
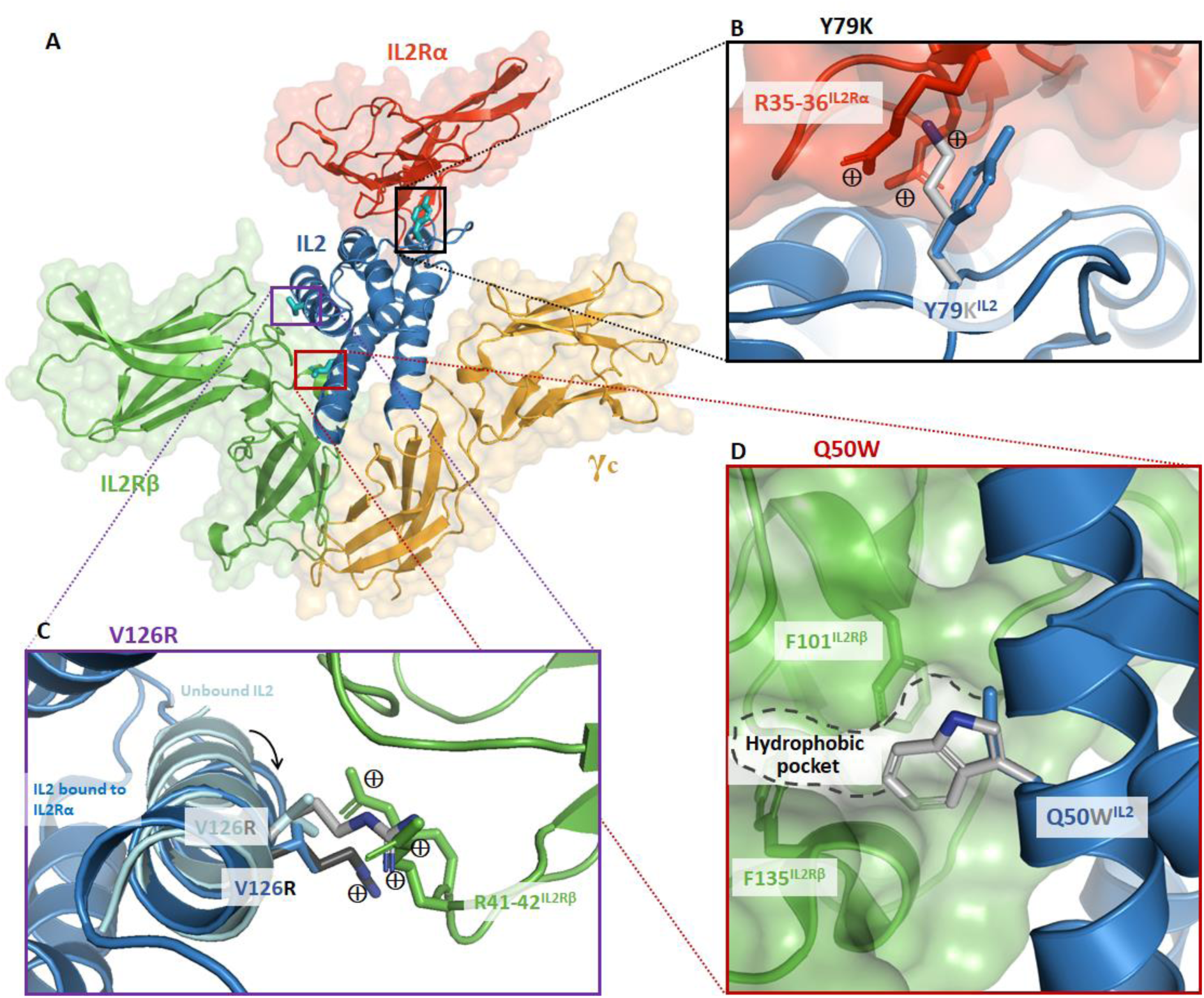
Structural modelling of murine IL2 muteins with the trimeric IL2R complex. **A)** Structure of IL2 in complex with the IL2RA/IL2RB/IL2RG trimeric complex, indicating the sites of the three mutated residues. **B)** The Y79K IL2 mutation, with close proximity to the R25-R26 residues in IL2RA. **C)** The V126R IL2 mutation, in relation to the R41-R42 residues in IL2RB. The structure of IL2 is shown based on both the unbound IL2 structure and the conformation of IL2 present in the IL2-IL2RA complex, indicating the rotation of the helix from which the V126R side-chain emerges. **D)** The Q50W IL2 mutation, in relation to the hydrophobic groove present between the F101 and F134 residues of IL2RB. Hydrophobicity in IL2RB represented by colour, with increasing intensity of green representing increasingly hydrophilic values.

### In vivo functional testing of V126R IL2 mutein

As proof-of-principle for the utility of the developed IL2 muteins for *in vivo* use, we sought to test the Treg-promoting V126R IL2 mutein in mice. We used an AAV-based system (AAV6.2 capsid and a club cell-specific promoter) for transgene delivery into the lung epithelium, (**Figure 4A**). Transduced cells, expressing the appropriate transcription factors to activate the delivered *Scgb1a1* promoter, are able to drive the production of IL2 in the lung tissue, without giving systemic IL2 production (manuscript in preparation). The system results in rapid production of IL2, with free IL2 levels dropping as IL2-consuming cells build up (**Figure 4B**). The V126R IL2 mutein built up at higher initial levels than the wildtype IL2 (**Figure 4B**), consistent with a limited number of IL2 consuming cells. The expression of wildtype IL2 in the lung increased the number of lung Tregs, while Tregs in other tissues remained normal (**Figure 4C**). The V126R IL2 mutein demonstrated a similarly restricted expression profile, producing an elevation in Treg numbers compared to wildtype IL2 (**Figure 4C**). This was achieved without expansion of the resident CD8 T cell population (**Figure 4D**), demonstrating improved *in vivo* properties of the V126R IL2 mutant in Treg expansion.

**Figure 4.**
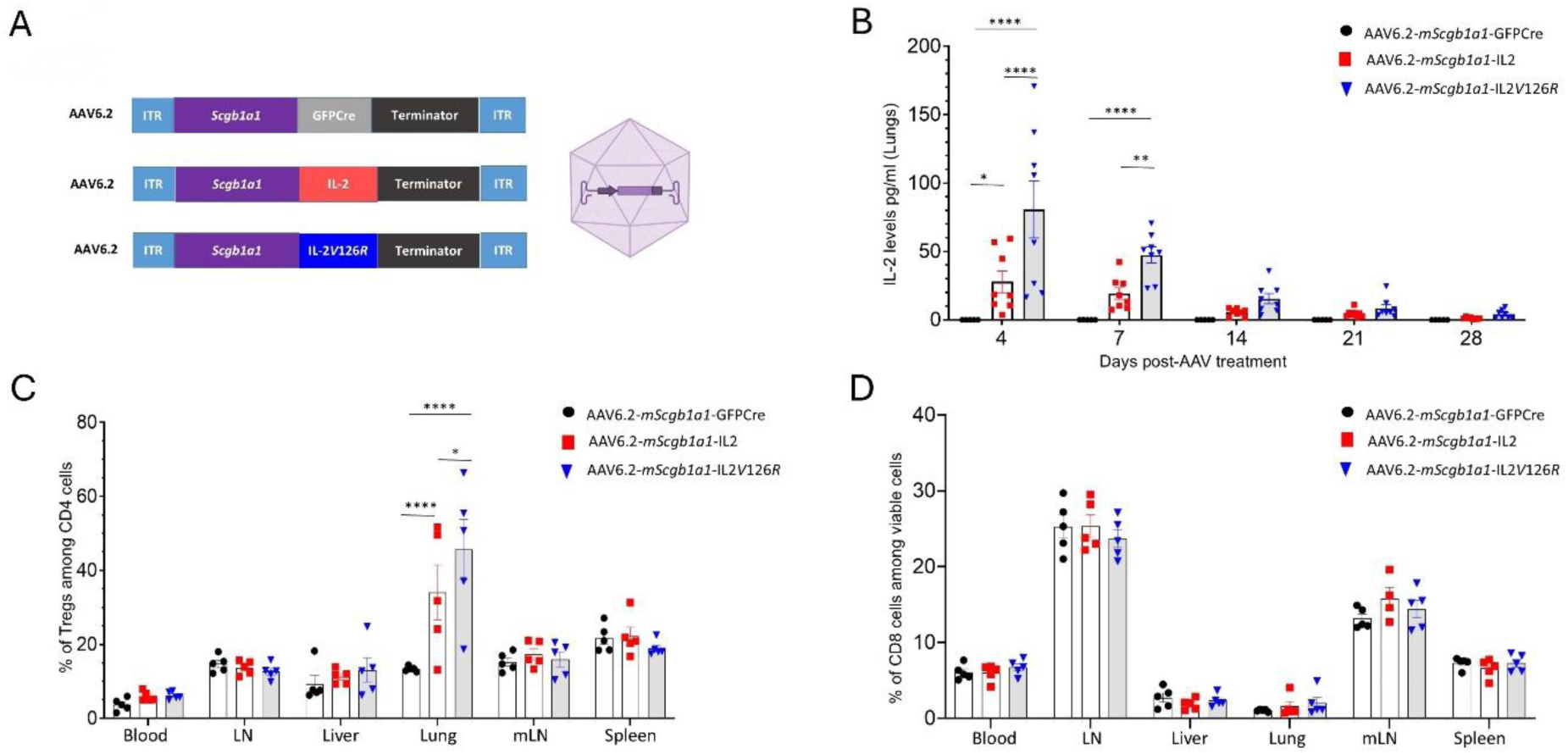
*In vivo* testing of V126R IL2 enhances local Treg expansion. IL2 and IL2 mutein were delivered *in vivo* using AAV-mediated gene delivery. **A.** Schematic representation of the viral vector used. Expression is driven by the *Scgb1a1* (Secretoglobin 1A1) promoter and flanked by the inverted terminal repeats. The expression cassette was contained within an AAV6.2 capsid which displays enhanced trophism for lung epithelial cells: AAVs were administrated through the intranasal route (n=5-7 mice/group). **B.** Available IL2 protein levels in the lung on days 4, 7, 14, 21 and 28 post-AAV delivery. **C.** Tissues were removed on day 14 and assessed via flow cytometry for the frequency of Foxp3+ Tregs within the CD4+ T cell population, and **D.** CD8 T cells within the viable cell population. All the data were analysed using Two Way Anova.

### Combinatorial generation of a CD8 T cell superkine

The muteins Q50W and Y79K both demonstrated different desirable properties as a CD8 T cell-promoting IL2 mutant. Q50W increased CD8 T cell signalling, while Y79K boosted production and impeded Treg signalling (**Figure 2C,D**). To determine if we could engineer a CD8 T cell superkine with all three properties, we sought to combine these mutations. Combination of Q50W Y79K rescued the mild production defect of Q50W, giving a similar production boost to that achieved by Y79K alone (**Figure 1A**. Upon titration, the Q50W Y79K mutant showed decreased Treg responses by ∼30 fold compared to both wildtype and Q50W alone (**Figure 5A**), giving a defect intermediate to that observed with Y79K. In CD8 T cells, responses to Q50W Y79K, mimicked that of Q50W (**Figure 5B**). The combination of Q50W Y79K therefore creates a cytokine with the ideal properties of the two parental mutants, with improved production, enhanced CD8 T cell responsiveness and impeded Treg responsiveness.

**Figure 5.**
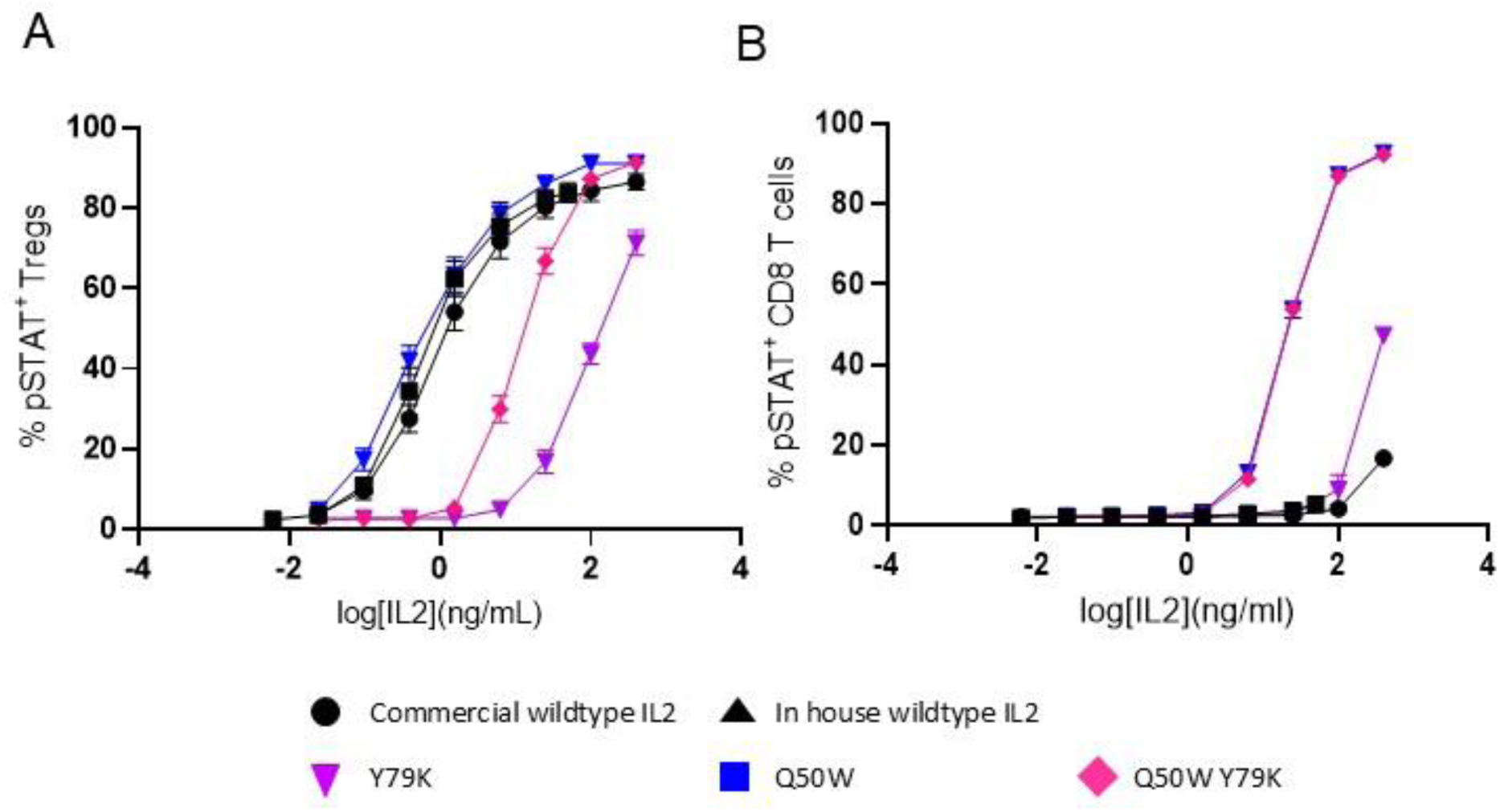
Combining mutations Q50W and Y79K enhances CD8 T cell selectivity of murine IL2. **A)** Mouse splenocytes were incubated for 20 minutes with either media containing IL2 or murine mutein, using a serial dilution of IL2 concentration, prior to staining for pSTAT5. The percentage of pSTAT5^+^ cells within the Treg or **B)** CD8 T cell populations (n=3).

### SolubiS human IL2 muteins have altered cellular specificity

Similar to the murine muteins, to determine the impact of the human IL2 mutations on receptor specificity, we first used a functional assay based on detection of signalling within human Tregs or CD8 T cells in mixed PBMCs. Of the mutations designed to prevent IL2 binding to the IL2RB (H36W, D40R, N108H, V111R), two of the mutations (D40R, N108H) performed poorly, with detrimental impacts on the pSTAT5 response in Tregs (**Figure 6A**). This suggests these muteins have a net defect in bioactivity due to poor recruitment of IL2RB to the IL2/IL2RA complex, potentially exacerbated by poor production (D40R=70.83 ng/ml, N108H 7.65 ng/ml). Three muteins, H36W, V111R and N108H T151R, by contrast, achieved the desired properties of maintained near-normal pSTAT5 responsiveness in Tregs, while also having a sharply reduced signal in CD8 T cells (**Figure 6B**). Notably, the V111R mutein is the human homolog of V126R, for which the structural modelling suggests IL2/IL2RA complex binding may be rescued by a conformational shift in the mutein residue (**Figure 3C**). For the residues designed to boost production, N110R, T143N and T151R had neutral effects on Treg signalling, although T143N was detrimental when combined with mutations designed to impede IL2RB binding (**Figure 6A**). The H36W, V111R and N108H T151R muteins were therefore candidates as Treg-promoting muteins, with poor IL2RB signalling, with V111R having marginally the best production of the three. This production was improved above wildtype values through *ad hoc* combination with N110R T151R, the SolubiS mutation with the best initial production values (**Figure 1B**).

**Figure 6.**
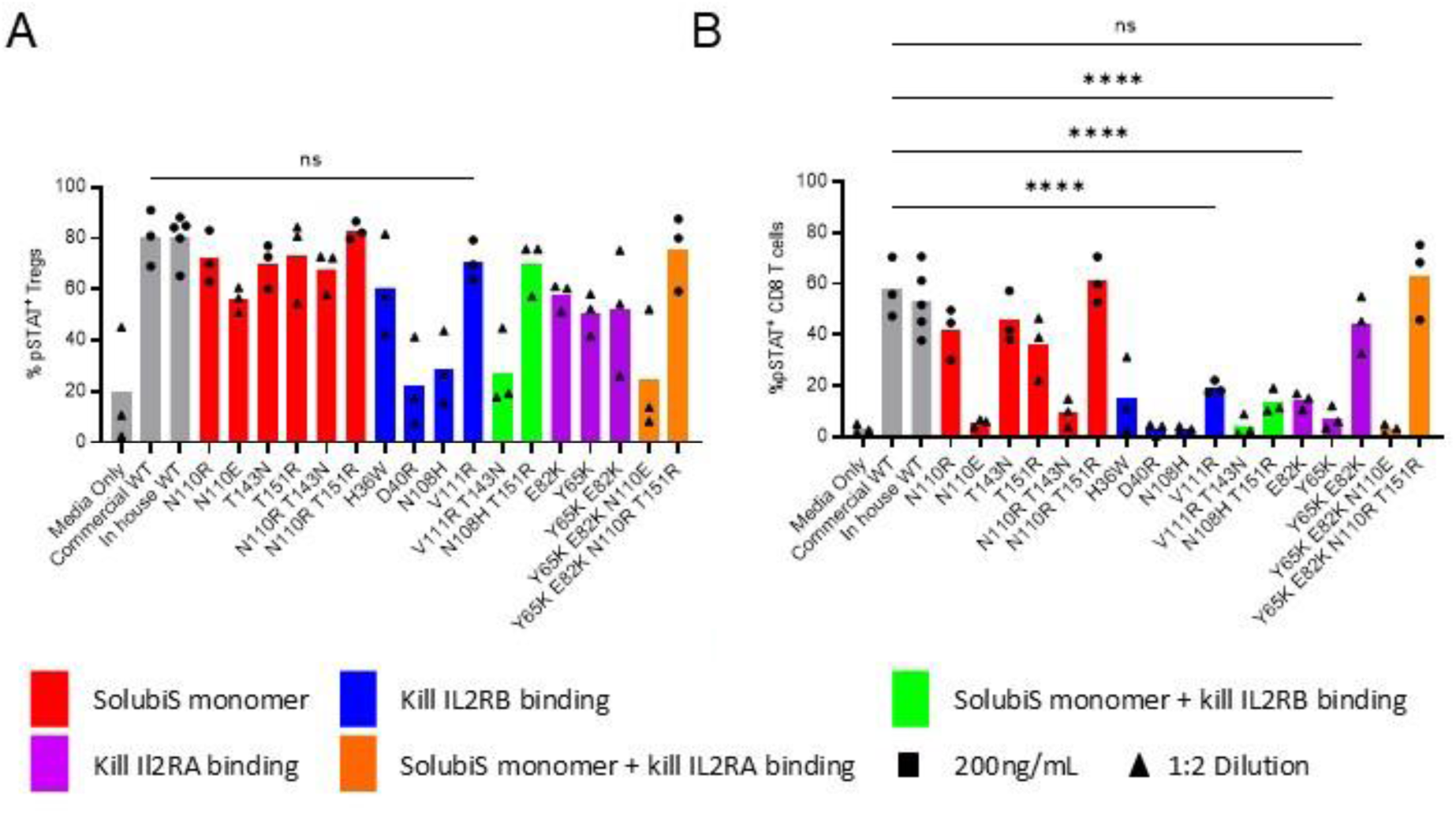
Altered cellular responses to human SolubiS IL2 muteins. **A)** Human PBMCs were incubated for 20 minutes with either media containing 200ng/mL human mutein or 1:2 dilution (where ELISA detected less than 200ng/ml), prior to staining for pSTAT5. The percentage of pSTAT5+ cells within the Treg or **B)** CD8 T cell populations (n=3). Comparisons made using One-Way ANOVA (*P < 0.05, **P < 0.01, ***P < 0.001, **** P<0.0001).

Of the muteins designed to prevent binding to IL2RA (E82K, Y65K, Y65K E82K) all three modestly reduced Treg responses as designed (**Figure 6A**). However, E82K and Y65K also did not produce pSTAT5 responses in CD8 T cells, potentially due to production issues lowering their concentration in the assay (E82K = 10.9 ng/ml, Y65K = 9.5 ng/ml) (**Figure 6B**), making them less useful as candidate cytokines. Interestingly, the combined Y65K E82K mutein stimulated marked responses in CD8 T cells despite production issues (used at a concentration 6-fold less than wildtype, 30.4 ng/ml) (**Figure 6B**). The Y65K E82K mutation therefore met the biological requirements for a CD8 T cell-promoting mutein, although its relatively poor production would impede therapeutic use. Fortunately, we found that while N110R T151R in isolation demonstrated no increase in production, when in combination with Y65K E82K it rescued the marked defect in production caused by the Y65K E82K mutation and so it was decided to take forward Y65K E82K N110R T151R (**Figure 1B**).

### Combination mutants result in human IL2 muteins with enhanced cellular selectivity

Based on this biological characterisation, four human muteins were of interest for further testing: V111R of particular interest for potential Treg specificity, and the V111R N110R T151R combination with boosted production; Y65K E82K, of particular interest for potential CD8 T cell specificity, and the Y65K E82K N110R T151R production-boosted version. We therefore characterised these four cytokines in the PBMC pSTAT assay for titrated responses to Treg and CD8 T cell activation.

For the potential Treg-boosting V111R and V111R N110R T151R muteins, a titration of the muteins on human PBMCs had intact pSTAT responses in Tregs down to 10ng/ml, with a drop-off compared to wildtype IL2 at lower concentrations (**Figure 7A**). In contrast, CD8 T cell responses were muted ∼100-fold, even at the highest doses tested (**Figure 7B**). These results confirm that the V111R mutein, like the analogous V126R mouse mutein, has enhanced specificity for Tregs while maintaining near normal bioactivity.

**Figure 7.**
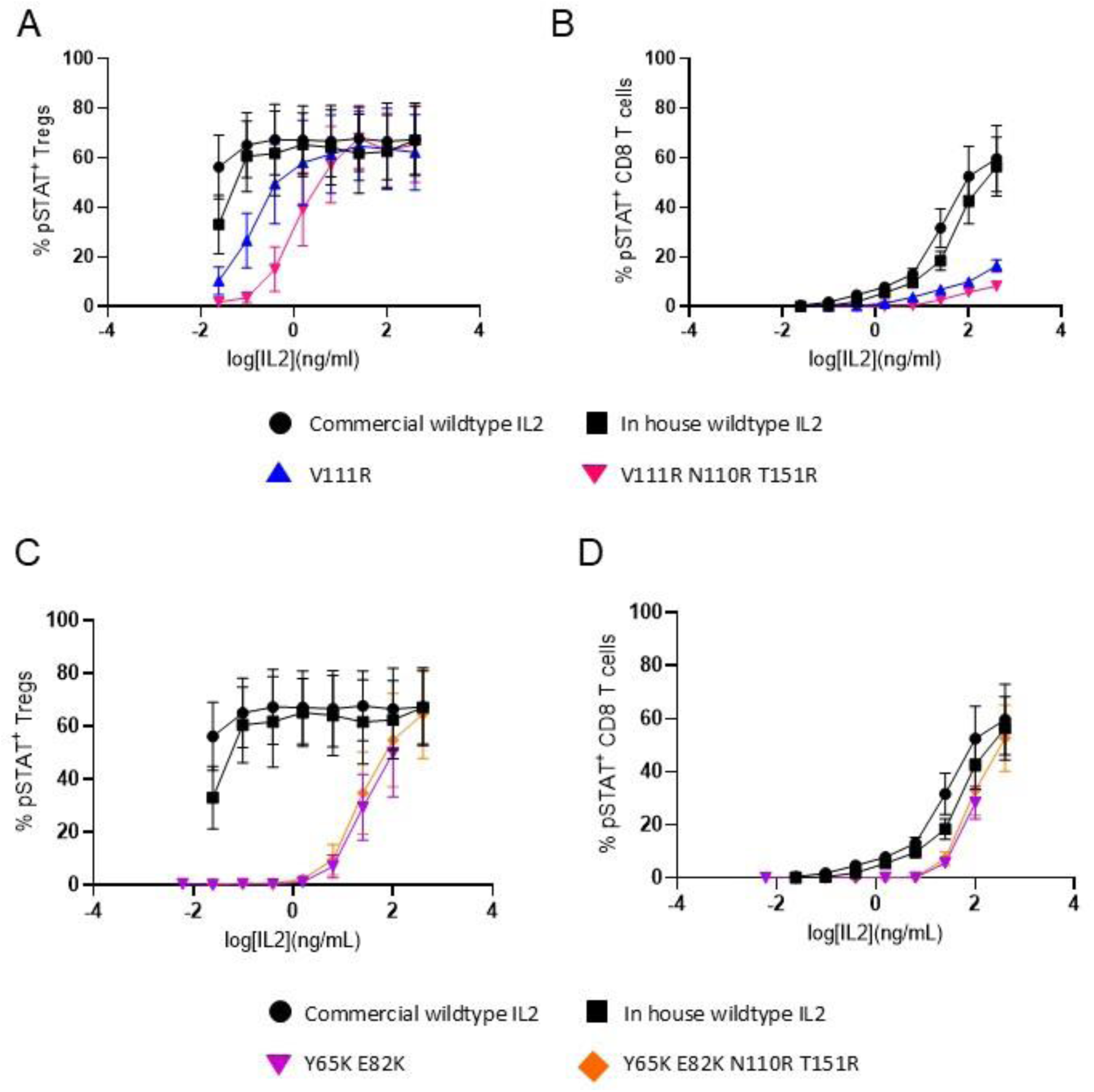
Cellular selectivity of Human IL2 muteins is maintained with addition of production boosting mutations N110R and T151R. Human PBMCs were incubated for 20 minutes with either media containing IL2 or human Treg selective (A,B) or CD8 T cell selective (C,D) mutein, using a serial dilution of IL2 concentration, prior to staining for pSTAT5. **A,C)** The percentage of pSTAT5^+^ cells within the Treg or **B,D)** CD8 T cell populations (n=3). All muteins were tested in the same experiment using the same controls.

For the potential CD8 T cell-selective muteins, Y65K E82K and Y65K E82K N110R T151R, production runs were tested in the same PBMC pSTAT assay. In titrations, both the Y65K E82K and Y65K E82K N110R T151R muteins showed sharply diminished Treg responses (∼3000 fold) (**Figure 7C**). In contrast, CD8 T cell responses were akin to wildtype (**Figure 7D)**, demonstrating strong CD8 T cell selectivity. The response curves for Treg and CD8 T cells were highly similar (**Figure 7C,D**), suggesting near-complete elimination of IL2RA binding and activation of Tregs only through the IL2RB-IL2RG complex, as in CD8 T cells. When also considering the enhanced production values of this mutein (**Figure 1B)**, overall, it shows a superior profile in CD8 T cell-stimulation.

## Discussion

Protein solubility is adapted to endogenous protein abundance in the cell, where protein folding occurs within the context of specific chaperones and potential interaction partners. For use in biotechnology or therapeutics, proteins require production in recombinant incubators, typically in cell types distinct from those in which they are naturally produced and frequently at concentrations that are several orders of magnitude above their physiological concentration. This often results in protein aggregation, reducing the total protein production yield and increasing costs, creating barriers in affordability and therefore availability of these drugs for patients. Aggregation is a complex process, influenced by physicochemical parameters such as protein and ion concentrations, pH, and temperature, making it challenging to identify sequences that will aggregate under native conditions. The solution to this challenge lies in the distinction between aggregation-prone regions that are thermodynamically protected by folding and those that occur in aggregation-competent conformations that can form without major unfolding transitions.

Using the SolubiS method, human IL2 was identified to have two aggregation-prone regions, while the mouse IL2 only has one. The first aggregation-prone region in human IL2 can be broken by the N110R or N110E mutations. Interestingly, the N110R solution identified by SolubiS is the naturally-occurrent residue present in mouse IL2 (R125), indicating that the mouse has found the same gatekeeper solution of an arginine side-chain in this location. The second aggregation-prone region was predicted to be improved by T143N or T151R (human) or A158N or T166R (mouse). T166R showed production improvements in the mouse as a solo mutation. A158N improved production when in combination with V126R, while T166R was required to restore the production efficiency of N123H. In the human muteins, the N110E residue was detrimental to both production and functional activity, either in isolation or in combination with specificity-enhancing mutations. While the N110R, T151R and T143N mutations did not boost production alone, they helped boost production of the impaired Y65K, E82K and V111R proteins. In addition to the cost savings advantages that the aggregation-protecting mutations could deliver for IL2 production (a therapeutic protein with annual sales of US$60 million), the results here demonstrate the extension of the SolubiS method for improving recombinant cytokine production[51], [49], an area of growing therapeutic importance.

Beyond the biotechnology value of increasing protein production, here we used the SolubiS method to simultaneously improve both the production level and the functional properties of the native IL2 protein, through the introduction of single amino acid changes. This functional improvement is theoretically easier for the mutations designed to enhance specificity for CD8 T cells. As CD8 T cells generally do not express high levels of IL2RA, blocking IL2RA binding can substantially reduce Treg responses, while having negligible impact on CD8 T cell responses. Indeed, we have identified Y65K E82K (human) and Y79K (mouse) muteins which have simple cation-cation repulsion against the IL2RA, while maintaining signalling through IL2RB/IL2RG complexes. In addition, we have identified a murine CD8 T cell superkine Q50W with enhanced IL2RB binding, which, when combined with Y79K becomes a CD8 T cell superkine with greater cellular selectivity. By contrast, muteins to enhance specificity for Tregs face a potential “functionality tax”. While IL2 first binds IL2RA, the recruitment of IL2RB into the complex enhances signalling through the trimeric receptor. Thus, any mutations which completely block IL2RB binding will kill CD8 T cell responses, but also dampen down bioactivity on Tregs. This “functionality tax” was observed on all Treg muteins, with the exception of V111R (human) / V126R (mouse). The success of these muteins likely lies in the structure of IL2 being nearly identical whether free or bound to the receptor, with the exception of the beginning of helix C. Following binding to IL2RA, the beginning of helix C is slightly unwound to move forward by 1.0 Å, a confirmation change thought to prime superior binding to IL2RB via a hydrogen bond formed between N88 of IL2 and R42 of IL2RB [39]. In its unbound state, the IL2 muteins V126R (mouse) and V111R (human) exhibit a cation-cation repulsion to R41-R42, which blocks IL2RB binding. However, the binding of IL2 to IL2RA shifts the V126R / V111R residues back, enabling recruitment of IL2RB. This conformational-dependency of the V126R / V111R mutein effect may explain why interaction with CD8 T cells, via IL2RB, results in ∼1000-fold reduction in signalling activity, whereas Tregs, which first require engagement of IL2RA, are still able to recruit IL2RB for full signalling capacity. This advantage of specificity without loss of bioactivity is unlike that observed in some other Treg-specific muteins, where the loss of IL2RB binding is accompanied by an aggregate defect in signalling capacity even in Tregs [26],[34], [35], [36]. The uniqueness of this solution also allows its potential combination with other mutations that directly enhance IL2RA binding [23], potentially creating a Treg superkine. Notably, V111R (when annotated as V91R) has been previously described as an IL2 antagonist [25]. We did not investigate competitive binding of V111R and wildtype IL2, but based on lack of signal in CD8 T cells, if V111R still binds IL2RB/IL2RG in a non-signalling conformation, it could potentially act as an antagonist in CD8 T cells, while acting as a bioactive signaller in Tregs.

IL2 (marketed as Aldesleukin) was first approved for use in humans in in 1992 using high doses to trigger CD8 T cell responses to aid tumour clearance in the treatment of renal carcinoma [52]. The short half-life of IL2, and low affinity for the IL2RB-IL2RG receptor, means large bolus doses were needed to reach the concentration needed to stimulate CD8 T cells. This resulted in severe side effects, such as capillary leak syndrome, being common in the initial clinical trials [53], and tumour clearance efficacy inferior to that later observed with immune checkpoint blockade. Since this early clinical use, the conceptual shift in our understanding of IL2 biology warranted re-orientation of the drug for anti-inflammatory use. As such, clinical trials have been run using multiple low-doses of IL2 to successfully at elevate Treg numbers in inflammatory conditions such as Graft Versus Host Disease and Systemic Lupus Erythematosus amongst others [54], [55], [56], [57]. Although trials have shown success in reducing disease symptoms, dosing remains a challenge due to the small therapeutic window. The muteins discussed here may help fix many of the caveats of IL2 therapy. Firstly, increased production efficiency of IL2 will help reduce costs making therapies more accessible to patients. Secondly increased specificity of the muteins will help ameliorate some of the side effects. It can be hypothesised that due to the selectivity of muteins Y65K E82K and Y65K E82K N110R T151R for CD8 T cells, lower doses may stimulate effective CD8 T cell responses, while simultaneously reducing the risk of severe side effects and increasing protein half-life due to less consumption by off-target Tregs. Likewise, but with an inverted hierarchy of specificity, the V111R or V111R N110R T151R muteins, with enhanced selectivity for Tregs, may further decrease the dose required to elicit therapeutic Treg responses. By combining selective IL2 muteins like those discussed here with novel delivery systems such as AAVs, which allow stable, tissue specific protein production [19], we can address many of the limiting characteristics of IL2 therapy, such as half-life, therapeutic window and off-target responses. Together these approaches have the potential to further enhance the efficacy of IL2 in the clinic for both anti- and pro-inflammatory uses.

## Supporting information

Supplementary spreadsheet 1. Predicted properties of designed mutations.

## Acknowledgements

The work was supported by the VIB, an ERC Proof of Concept Grant TreatBrainDamage (to A.L.), the Biotechnology and Biological Sciences Research Council through Institute Strategic Program Grant funding BBS/E/B/000C0427 and BBS/E/B/000C0428, and the Biotechnology and Biological Sciences Research Council Core Capability Grant to the Babraham Institute. MGH is currently the ERANet Chair (NCBio) at i3S Porto funded by the European Commission (H2020-WIDESPREAD-2018-2020-6; NCBio; 951923). The authors acknowledge the important contributions of the Babraham Institute Flow Cytometry Core and Biological Services Unit and the University of Cambridge FACS Core and University Biological Services Unit.

## Authors Contribution

A.D., N.M., A.G., A.H, F.N., M.A. S.L. and O.B. designed and performed experiments, and evaluated and interpreted data. R.vdK. performed in silico design and provided intellectual input. J.D., M.G.H., F.R., J.S., and A.L. conceptualized, supervised and financed the study. A.D. and A.L. drafted the initial manuscript and all authors contributed to the final version.

