## Supplementary Table 1 for "Directed disruption of IL2 aggregation sites produces designer biologics with enhanced specificity coupled to improved production capacity"

**Supplementary Table 1**: List of antibodies used for IL2 experiments *in vivo*

| Antigen | Fluorophore | Clone | Dilution | Vendor | Stage |
|---|---|---|---|---|---|
| Viability | eFluor780 | -- | 4000 | Thermofisher | Viability |
| Streptavidin | AF350 | -- | 500 | Invitrogen<br>Cat No. S11249 | Streptavidin |
| CD45 | BUV395 | 30-F11, rat IgG2b | 2000 | BD Biosciences<br>Cat No. 564279 | Surface |
| GATA-3 | eFluor450 | TWAJ, rat IgG2b | 500 | ThermoFisher<br>Cat No. 48-9966-42 | Intracellular |
| CD25 | BV480 | PC61, rat IgG1 | 500 | BD Biosciences<br>Cat No. 566120 | Surface |
| CD44 | BV510 | IM7, rat IgG2b | 2000 | BioLegend<br>Cat No. 103044 | Surface |
| ICOS | BV605 | C398.4A, armenian hamster | 500 | BioLegend<br>Cat No. 313538 | surface |
| RORgT | PE | B2D, rat IgG1 | 500 | ThermoFisher<br>Cat No. 12-6981-80 | Intracellular |
| T-bet | PE-Dazzle594 | 4B10, mouse IgG1 | 200 | Biolegend<br>Cat No. 644828 | Intracellular |
| CD8 | PE-Fire640 | 53-6.7, rat IgG2a | 5000 | Biolegend<br>Cat No. 100790 | Surface |
| CD69 | PE-Cy5 | H1.2F3, armenian hamster | 500 | BioLegend<br>Cat No. 104510 | Surface |
| CD62L | PE-Cy7 | MEL-14, rat IgG2a | 1000 | BioLegend<br>Cat No. 104418 | Surface |
| CD4 | APC-Fire810 | GK1.5, rat IgG2b | 500 | BioLegend<br>Cat No. 100480 | Surface |
| Foxp3 | APC | REA788, human IgG1<br>FJK-16s, rat IgG2a | 200; 200 | Miltenyi Biotec<br>Cat No. 130-111-601<br>ThermoFisher<br>Cat No. 17-5773-82 | Intracellular |
| mouse IgG | Unconjugated | Mouse IgG Isotype Control | 200 | Invitrogen<br>Cat No. 10400C | Blocking |
| rat IgG | Unconjugated | Rat IgG Isotype Control | 200 | Invitrogen<br>Cat No. 10700 | Blocking |
| 2.4G2 supernatant | -- | -- | | In-house | Blocking |
