## Supplementary Spreadsheet 1 for "Directed disruption of IL2 aggregation sites produces designer biologics with enhanced specificity coupled to improved production capacity"

**Supplementary spreadsheet 1. Predicted properties of designed mutations.** dTANGO, aggregation propensity score (reduction by >100 considered to reduce aggregation); ddG, stabilisation score (> +0.5 as destabilising, ddG < -0.5 as stabilising); ddGComplex, interaction energy (> +0.5 as destabilising, ddGcomplex < -0.5 as stabilising). ddG and ddGComplex given in relation to the structures 1m47 (IL2 alone), 1z92 (IL2 with IL2RA), 2b5i and 2erj (IL2 with IL2RA/IL2RB/IL2RB).
